# Invertebrate FAXC and Metaxin Proteins: Related Proteins with Distinct Features

**DOI:** 10.64898/2026.01.05.697642

**Authors:** Kenneth W. Adolph

## Abstract

Structural features of FAXC proteins of invertebrates are examined in this report to further establish that invertebrate FAXCs are metaxin-like proteins. Invertebrate phyla that receive the most attention are Mollusca, Cnidaria, and Arthropoda. The results demonstrate that invertebrate FAXC proteins have structural homology to vertebrate and invertebrate metaxin proteins. However, pairwise alignments of amino acid sequences revealed only low percentages of amino acid identities, indicating that FAXC and metaxin proteins are distinct categories of proteins. Among the important criteria in identifying invertebrate FAXC proteins as metaxin-like was the presence of conserved protein domains: GST_N_Metaxin, GST_C_Metaxin, and Tom37. Another metaxin-like feature found with invertebrate FAXC proteins is a characteristic pattern of α-helical and β-strand segments. The four β-strand segments form, in three dimensions, a planar β-sheet motif that is a distinguishing characteristic of FAXC and metaxin proteins. Evolutionary relationships between FAXCs and metaxins were uncovered by phylogenetic analysis. Distinct but related evolutionary groups were found for invertebrate FAXCs and metaxins, as well as vertebrate FAXCs and metaxins. Invertebrate species can have multiple FAXC genes, and more than ten FAXC genes is not uncommon. The genes that are adjacent to different FAXC genes in the same invertebrate are typically not the same. Therefore, the genomic regions are generally not highly conserved for multiple FAXC genes. But, for invertebrates with a single FAXC gene, the same neighboring genes may be found for the FAXC genes of different but taxonomically related invertebrates. Honey bees and bumblebees are examples. For invertebrates with multiple FAXC genes, several of the genes may be in close proximity in the same genomic region, forming a cluster of FAXC genes.

## 1. INTRODUCTION

Because very little is known about FAXC proteins and genes, this study was undertaken to examine in greater detail than previously the major structural features of FAXC proteins, specifically in invertebrates. The failed axon connections (*fax*) gene was first reported in the fruit fly *Drosophila melanogaster* (Hill et al., 1995). It was described as a novel *Drosophila* gene that is expressed as the Fax protein in axons of the central nervous system. No further publications concerning the *Drosophila fax* gene have appeared in the literature. However, searches of NCBI databases have shown that FAXC genes are widely distributed in a variety of vertebrates, invertebrates, plants, protists, fungi, and bacteria. For humans, the FAXC gene, also designated *C6Orf168* (Chromosome 6 Open Reading Frame 168), is located at 6q16.2. An initial investigation of vertebrate and invertebrate FAXC proteins, with amino acid sequences predicted from genomic sequences, revealed a strong similarity to metaxin proteins (Adolph, 2023). This report is to build upon these initial observations.

The primary structural features that are common to both FAXC proteins and metaxin proteins include GST_N_Metaxin and GST_C_Metaxin protein domains. In addition, metaxin and FAXC proteins possess the same highly conserved pattern of α-helical and β-strand segments. The β-strand segments form a special planar β-sheet motif produced by the four closely apposed β-strands. The motif is a distinguishing characteristic of FAXC and metaxin proteins. Further, phylogenetic analysis demonstrated that FAXC and metaxin proteins are related to each other by evolution.

Metaxin 1 was the first metaxin protein to be identified. It was demonstrated to be a component of the outer membrane of mitochondria in the mouse and in humans. A role for the protein has been implicated in the process of protein uptake into mitochondria. Metaxin 2, with only low homology to metaxin 1, was revealed to be a protein that associates with metaxin 1. A third metaxin, metaxin 3, was more recently identified (Adolph, 2019), and, like metaxins 1 and 2, has a wide occurrence among vertebrates. However, the role of metaxin 3 is not known. Many other organisms of different phyla have metaxin-like proteins. These include species of plants, protists, fungi, and bacteria.

Having a fully sequenced genome was an important criterion in selecting invertebrates to be included in this study. By providing information about the proteins encoded by genomic DNA, genome sequences are key elements in revealing similarities and differences in protein species and evolutionary relationships of organisms, including invertebrates. Also considered as criteria were the health, economic, and environmental significance of the invertebrates selected.

Molluscs in this study include the Mediterranean mussel *Mytilus galloprovincialis* (genome sequence: Gerdol et al., 2020) and the Pacific oyster *Crassostrea gigas* (Zhang et al., 2012), both economically important. The Cnidaria phylum is represented by the freshwater polyp *Hydra vulgaris* (Simakov et al., 2022), a model organism for regeneration research. Another cnidarian is *Stylophora pistillata* (Voolstra et al., 2017), a species of stony coral. The arthropod *Daphnia pulex* (Colbourne et al., 2011) is the most common species of water flea and the first crustacean to have its genome sequenced. A commercially important arthropod is the American lobster *Homarus americanus* (Polinski et al., 2021). Insects (phylum: Arthropoda; class: Insecta) in this investigation include the domestic silk moth *Bombyx mori* (Xia et al., 2008), the western honey bee *Apis mellifera* (Weinstock et al., 2006), which is the primary honey bee species, and the common eastern bumblebee *Bombus impatiens* (Sadd et al., 2015), one of the main species of pollinator bees. The Mediterranean fruit fly *Ceratitis capitata* (Papanicolaou et al., 2016), a destructive pest of fruit, is also part of the study. An invertebrate of the Chordata phylum is *Branchiostoma floridae* (Putnam et al., 2008), the Florida lancelet or amphioxus. Vertebrates used for comparison with invertebrates are, besides humans, the mouse (genome sequence: Mouse Genome Sequencing Consortium, 2002), frog (*Xenopus laevis*; Session et al., 2016), and zebrafish (*Danio rerio*; Howe et al., 2013).

The aim of this investigation was to examine in detail the structural features of invertebrate FAXC proteins. In particular, invertebrate FAXC proteins were compared to the metaxin proteins of invertebrates and vertebrates, since FAXC and metaxin proteins have structural features in common. The results confirm that FAXC proteins of invertebrates, like those of vertebrates, belong to the metaxin-like family of proteins. The findings extend the observations previously made for the FAXC proteins of vertebrates and invertebrates (Adolph, 2023). Structural features shared by both FAXC and metaxin proteins of invertebrates include GST_N_Metaxin and GST_C_Metaxin protein domains, similar patterns of α-helical secondary structure, and a special 3D β-sheet motif. The results presented in this report provide fundamental information about invertebrate FAXC proteins, and lay the groundwork for additional investigations into the structure and function of these proteins.

## 2. METHODS

Amino acid identities and positives in aligning two protein sequences were detected with the NCBI Global Align and Align Two Sequences tools (https://blast.ncbi.nlm.nih.gov/Blast.cgi; Needleman and Wunsch, 1970; Altschul et al., 1990). The examples in Figure 1 used Global Align, which compares the full lengths of two sequences. The major conserved protein domains (GST_N_Metaxin, GST_C_Metaxin, Tom37) of invertebrate FAXC proteins were revealed with the NCBI CD search tool, which is available at www.ncbi.nlm.nih.gov/Structure/cdd/wrpsb.cgi (Lu et al., 2020). Figure 2 includes the conserved domains for a variety of invertebrates. Identifying the α-helical and β-strand secondary structures of FAXC and metaxin proteins made use of the PSIPRED secondary structure prediction server. The server is found at bioinf.cs.ucl.ac.uk/psipred/ (Jones, 1999; Buchan and Jones, 2019). Multiple-sequence alignments were carried out with the COBALT multiple-sequence alignment tool at www.ncbi.nlm.nih.gov/tools/cobalt (Papadopoulos and Agarwala, 2007). COBALT was employed to compare the alignments of α-helical and β-strand secondary structure segments for different FAXC proteins (Figure 3). The 3D structures predicted for invertebrate FAXC proteins, and particularly the characteristic β-sheet motif (Figure 4), were investigated with the AlphaFold protein structure database (Jumper et al., 2021; Varadi et al., 2022) and the iCn3d structure viewer (Wang et al., 2020). Phylogenetic trees in Figure 5 were produced using the COBALT multiple-sequence alignment tool, with the phylogenetic trees generated from the alignments. The genes that are adjacent to invertebrate FAXC genes, as shown in Figure 6, were identified with the NCBI “Gene” database, specifically through “Genomic regions, transcripts, and products”. The “Gene” database was also the source of the genomic regions with multiple FAXC genes, such as in Figure 7.

**Figure 1.**
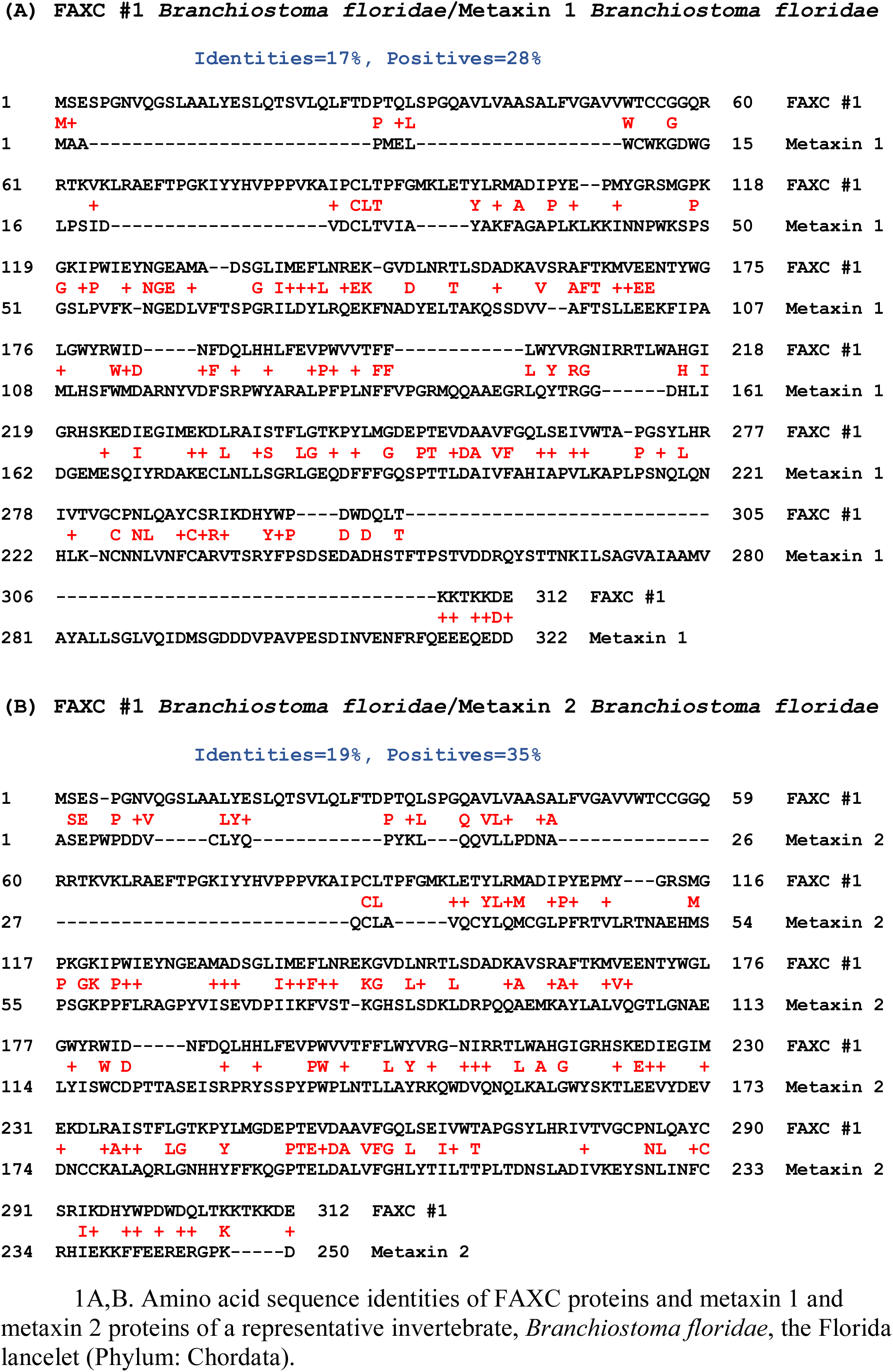
Amino acid sequence alignments of FAXC and metaxin proteins of invertebrates. In (A), FAXC #1 and metaxin 1 of *Branchiostoma floridae* are aligned. NCBI Global Align reveals only a low percentage of identical amino acids (17%). In (B), FAXC #1 and metaxin 2 of *B. floridae* are similarly aligned, with again a low percentage of identical amino acids (19%). The FAXC and metaxin proteins of different invertebrates show similar results. The conclusion can be drawn from these and other alignments that FAXC and metaxin proteins are different classes of proteins.

**Figure 2.**
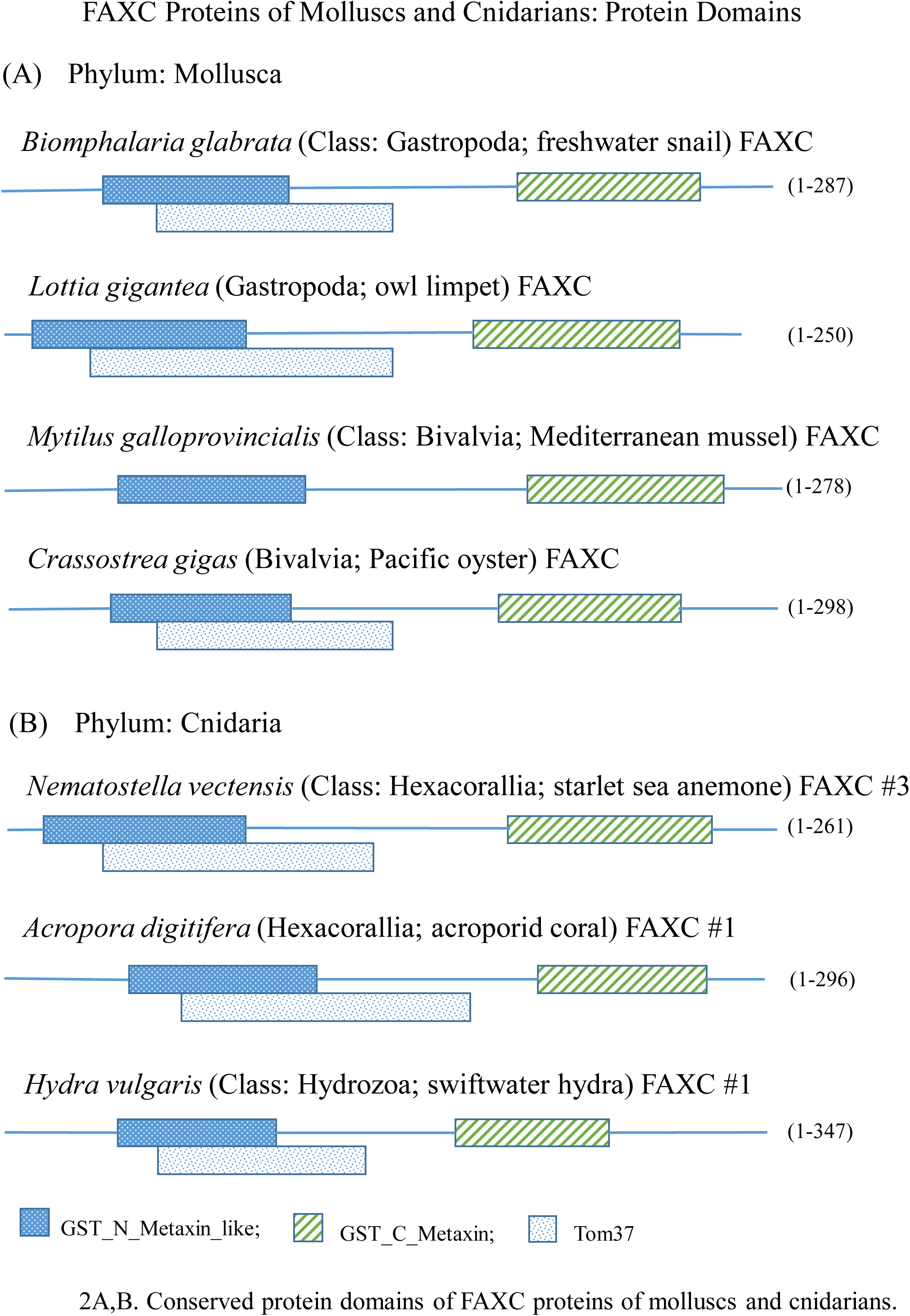
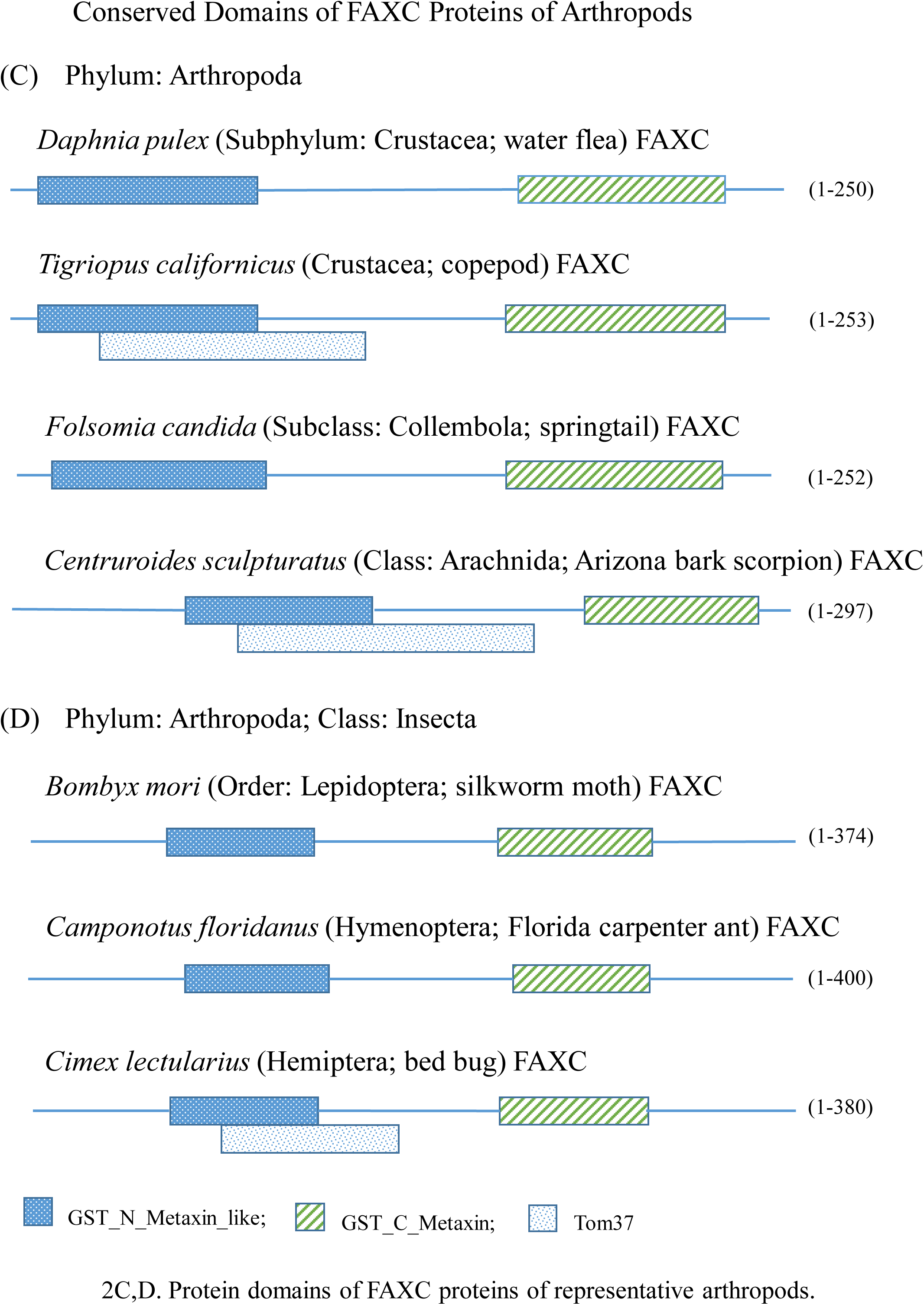
Major, characteristic protein domains of invertebrate FAXC proteins. The figure shows the primary protein domains (GST_N_Metaxin, GST_C_Metaxin, and Tom37) of FAXC proteins of representative invertebrates. The FAXCs are from invertebrate phyla that include Mollusca (Figure 2A), Cnidaria (2B), and Arthropoda (2C) with emphasis on Insecta (2D). All of the examples possess the GST_N_ and GST_C_Metaxin domains. The Tom37 domain is present in most of the FAXCs, though not in all. Invertebrates of phyla such as Cnidaria can have more than a single FAXC gene. For example, *Nematostella vectensis*, the starlet sea anemone, has at least 6 FAXC genes. In Figure 2B, FAXC #1 and FAXC #3 refer to the proteins encoded by FAXC genes #1 and #3.

**Figure 3.**
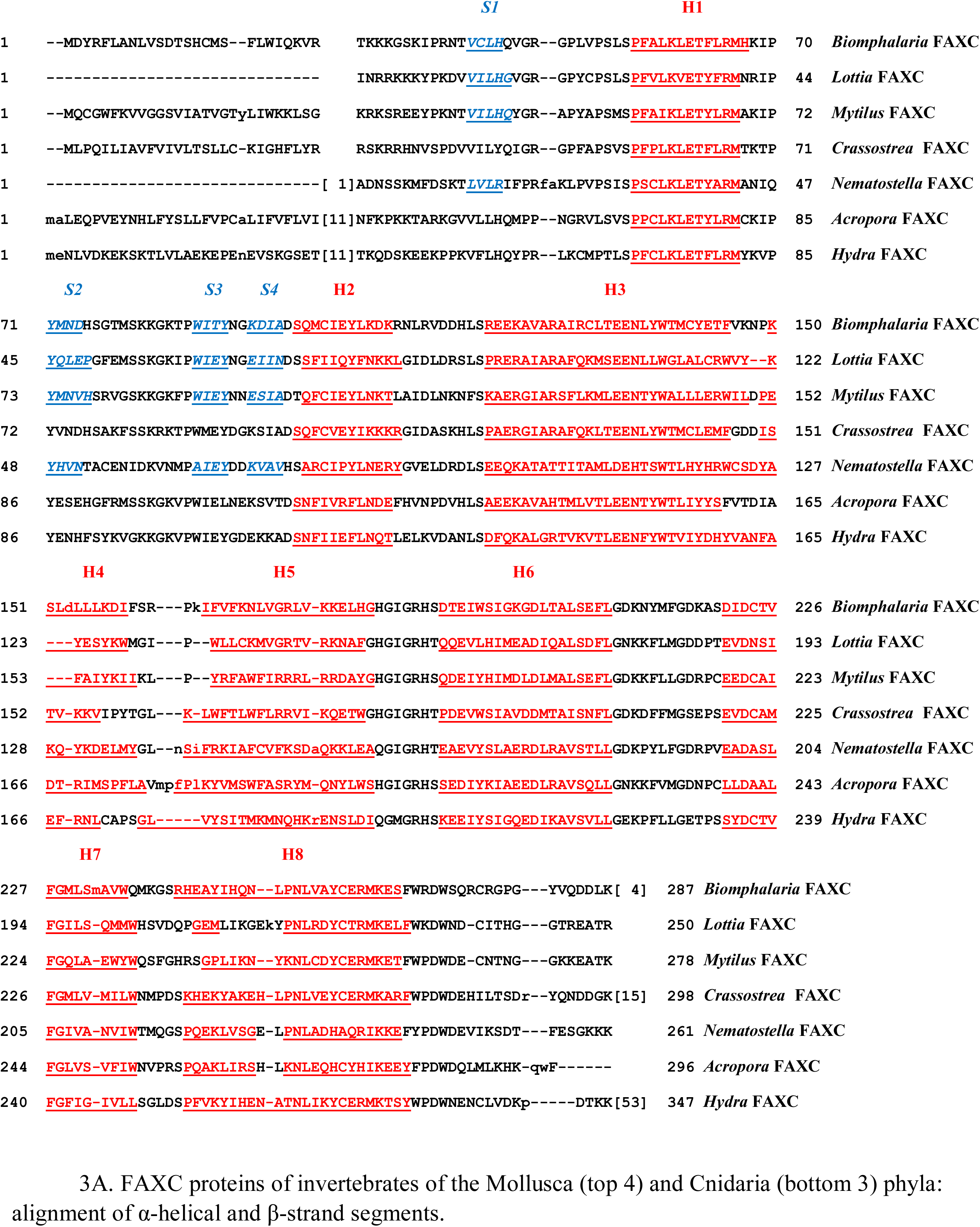
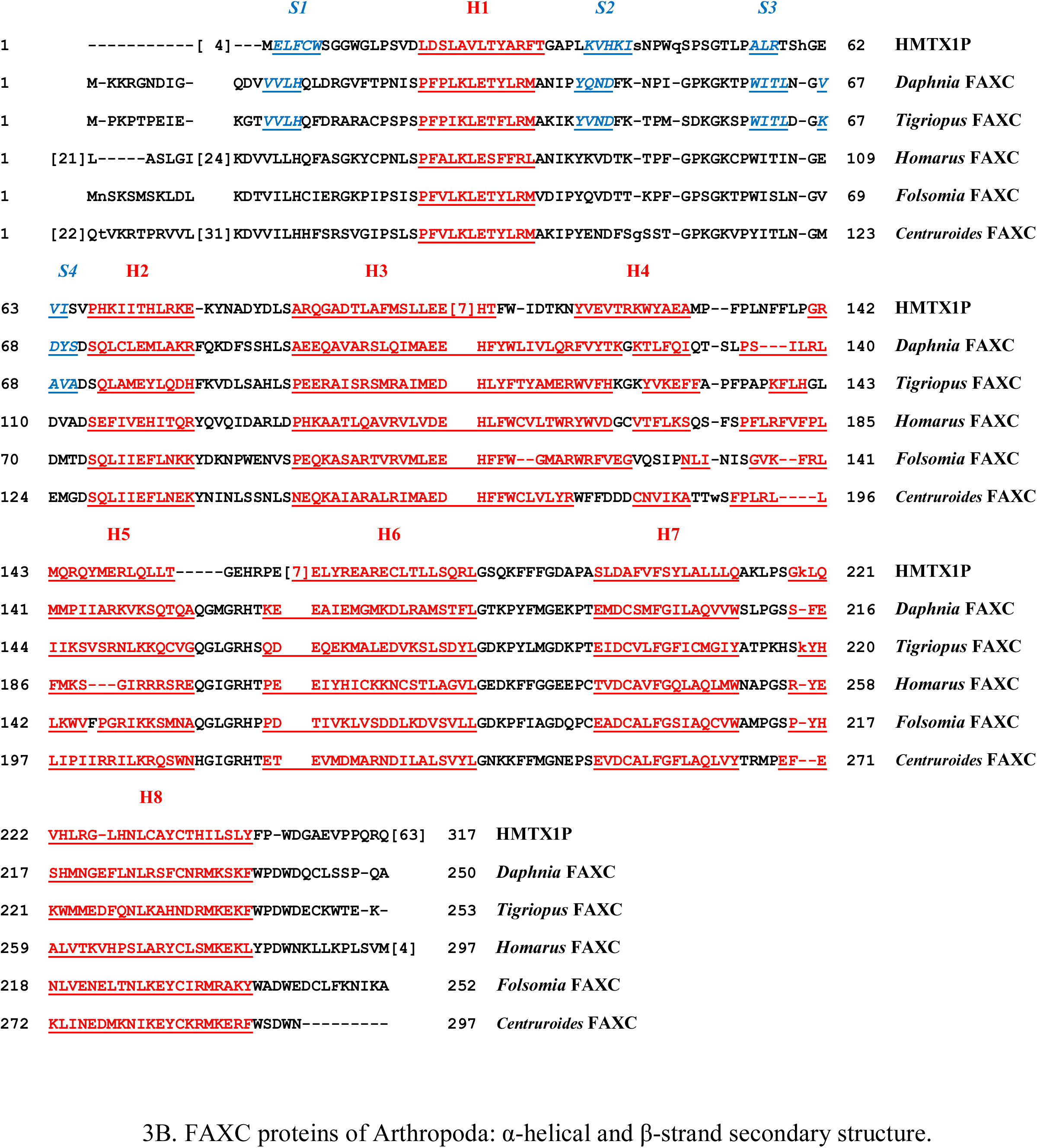
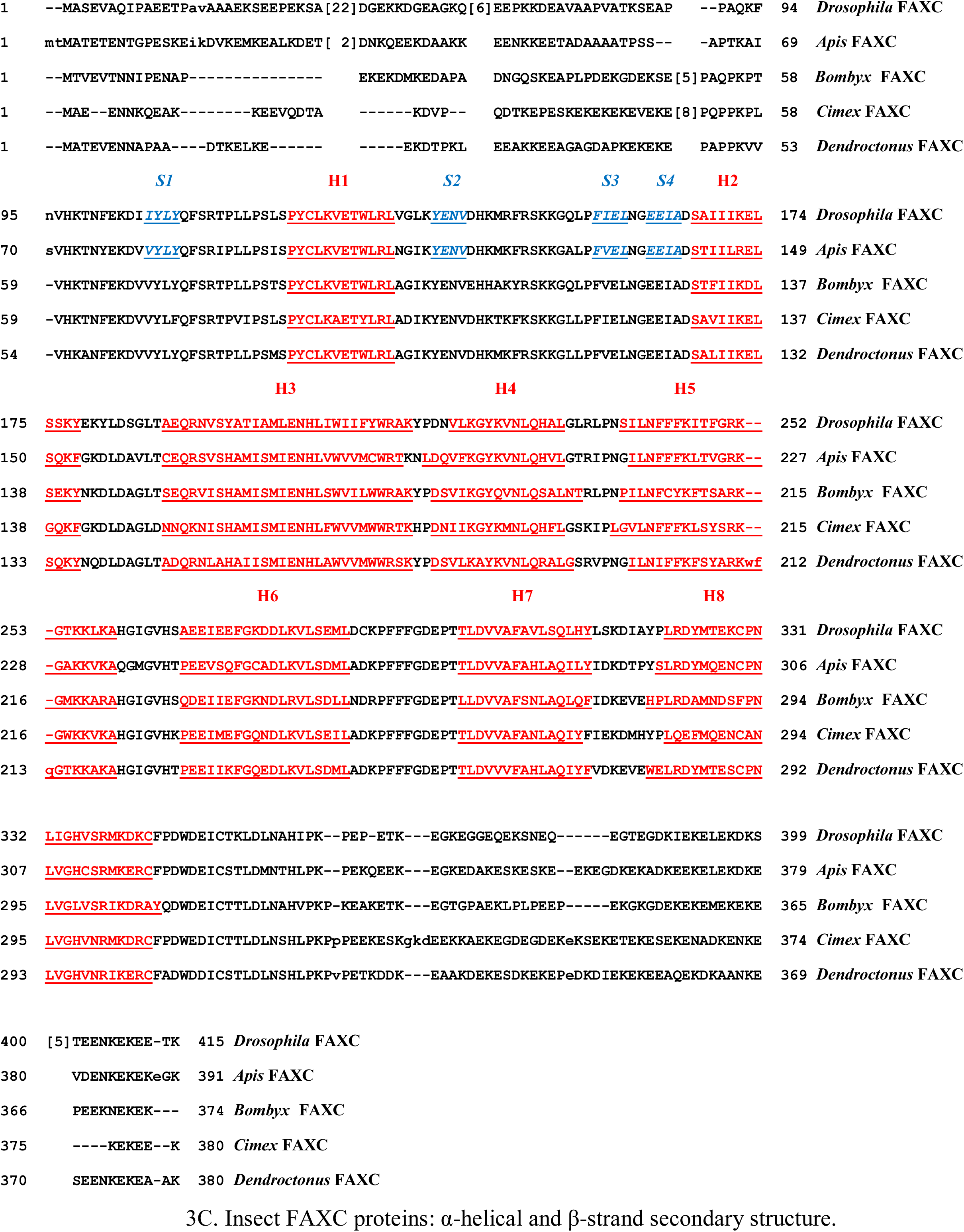
Conserved pattern of α-helical and β-strand secondary structures of invertebrate FAXC proteins. A highly conserved pattern of α-helical segments, H1 – H8 as shown in the figure, is a striking feature of invertebrate FAXC proteins. The amino acid sequences of the helical segments have a red color and are underlined. Figure 3A shows the protein secondary structures of representative molluscs (first four sequences) and cnidarians (bottom three sequences). Figure 3B includes the FAXC sequences of five arthropods, and also human metaxin 1 (HMTX1P). Five insect FAXC protein sequences are in Figure 3C (Phylum: Arthropoda; Class: Insecta). The patterns of α-helices in Figure 3A, 3B, and 3C are almost identical, emphasizing the characteristic and highly conserved nature of the H1 – H8 helical pattern in FAXCs and metaxins. Figure 3 also shows the β-strand segments of the invertebrate FAXC proteins. The β-strand sequences, and the examples in Figure 4 of the characteristic β-sheet motif, were obtained from the 3D protein structures predicted by AlphaFold (Jumper et al., 2021). The β-strand segments are those currently available for the invertebrate FAXCs in the figure. Like the α-helices, the β-strand segments form a highly conserved pattern. This consists of four, short β-strands, labeled S1 – S4 and shown with a blue color. The four β-strand segments were the only β-strands found for the proteins in Figure 3. Strand S1 precedes helix H1, while S2, S3, and S4 are between H1 and H2. The same pattern of β-strands is found for invertebrate phyla that include Mollusca (Figure 3A), Cnidaria (3A), Arthropoda (3B), and the Insecta class of Arthropoda (3C). The pattern of four β-strands is found not only for invertebrate FAXC proteins, but also for the metaxin proteins, such as human metaxin 1 (HMTX1P) in Figure 3B.

**Figure 4.**
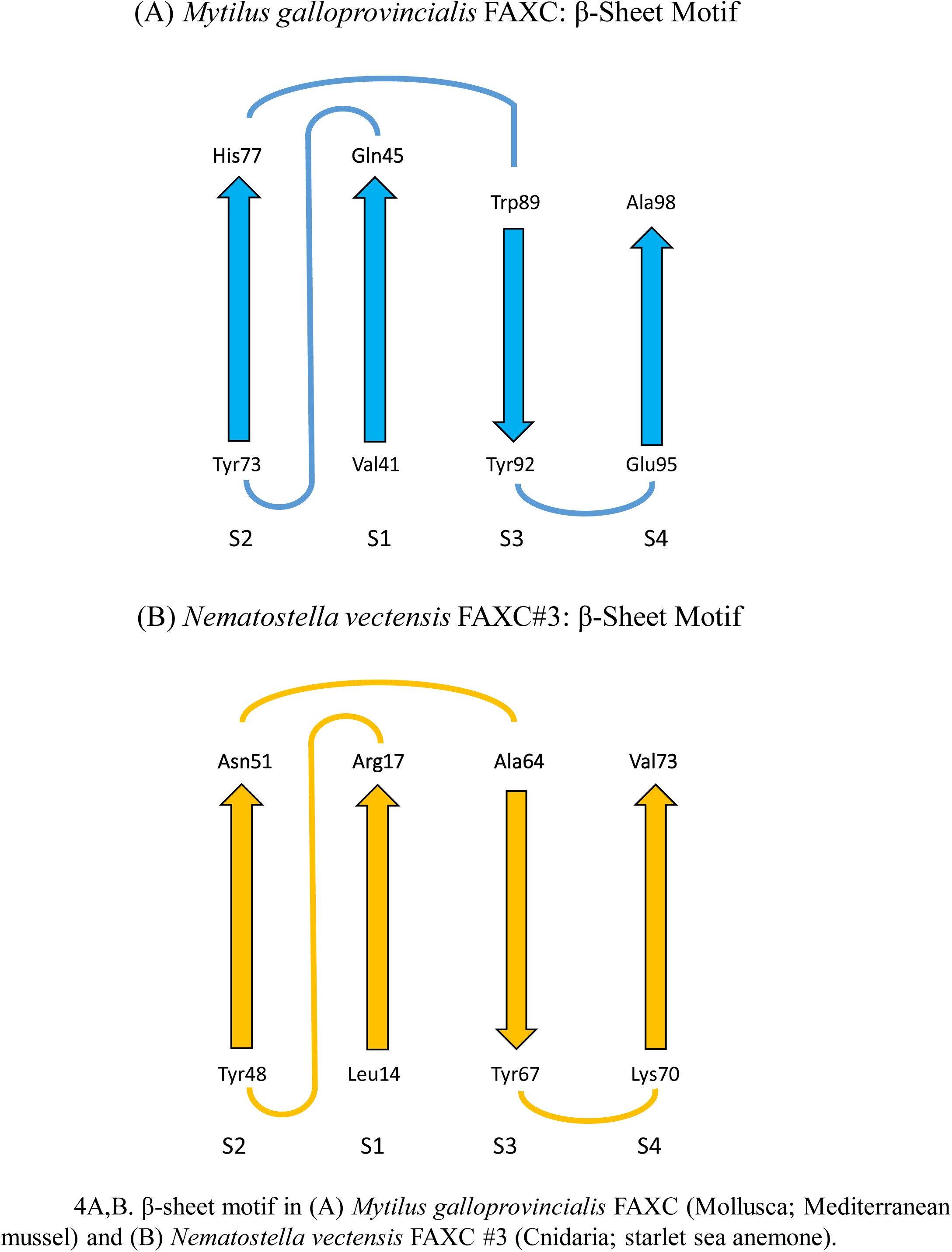
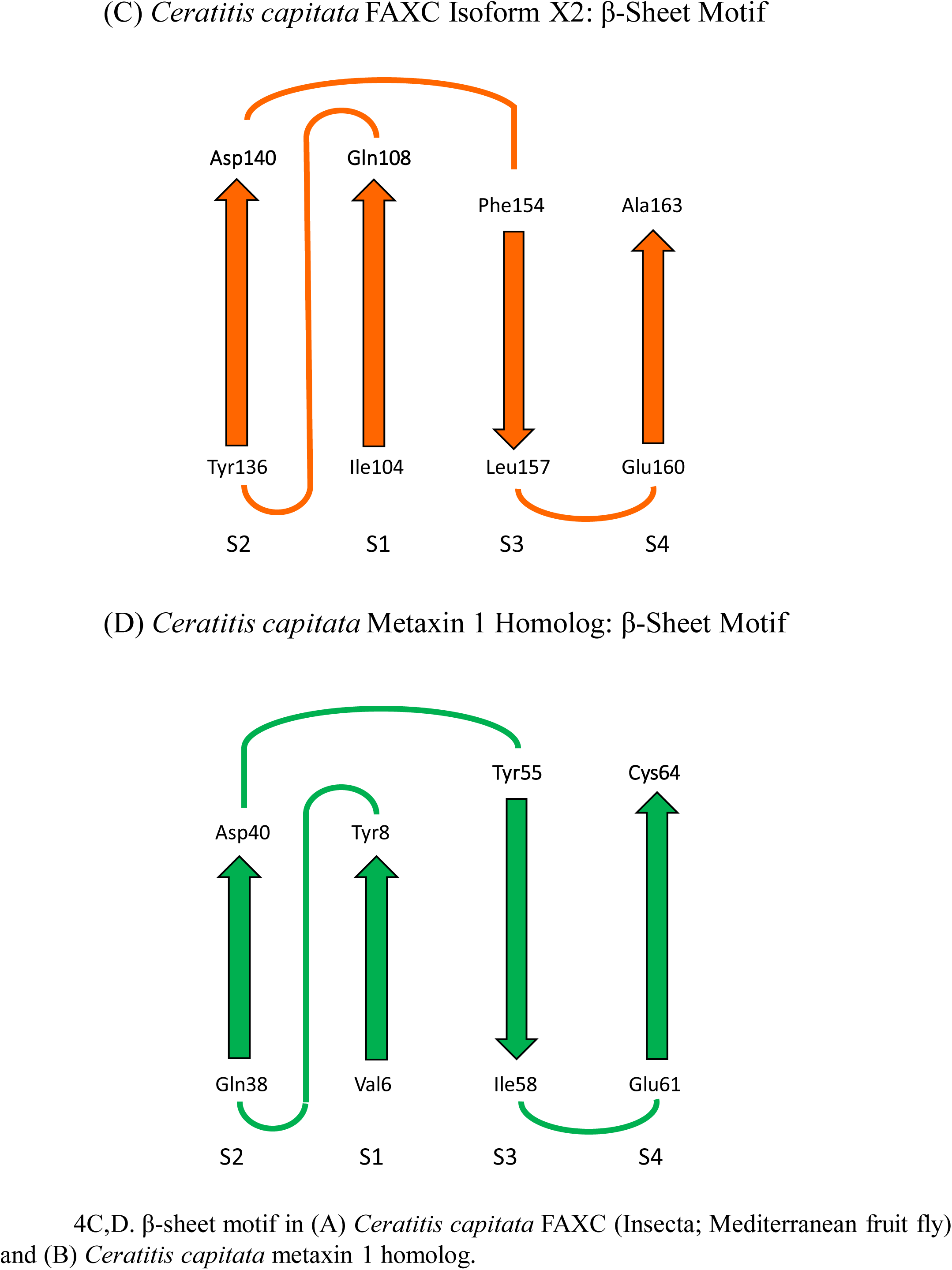
Characteristic β-sheet motif of invertebrate FAXC proteins. The predicted 3D structures of invertebrate FAXC proteins revealed that the β-strands in Figure 3 are arranged in three-dimensions as a four-stranded β-sheet. The β-sheet motif is highly conserved among invertebrate FAXC and metaxin proteins. Examples are shown in 4A, B, C, and D. Figure 4A includes the β-sheet structure in a representative mollusc, *Mytilus galloprovincialis*. An approximately planar structure is formed with the strands in a parallel and antiparallel arrangement. The β-sheet motif is typical of FAXC and metaxin proteins in consisting of four strands with the order and directions S2 (↑), S1 (↑), S3 (↓), S4 (↑). The β-sheet structure in *Nematostella vectensis*, a cnidarian (sea anemone), follows a similar arrangement (Figure 4B). In (C) and (D), the β-sheet structures in both the FAXC and metaxin proteins of the Mediterranean fruit fly *Ceratitis capitata* are compared. The structures are similar, strengthening the conclusion that the β-sheet motif is highly conserved and that FAXCs are metaxin-like proteins.

**Figure 5.**
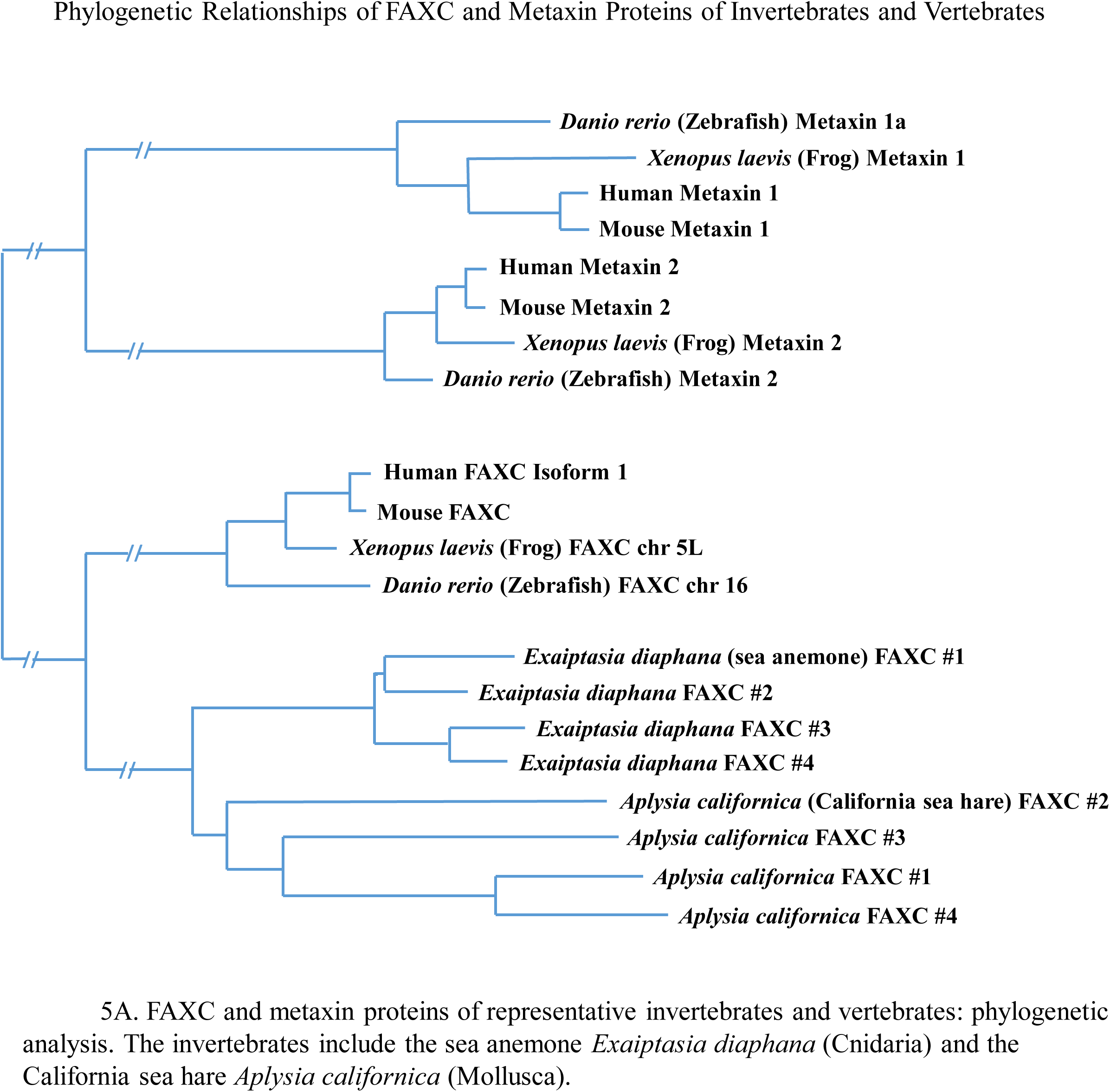
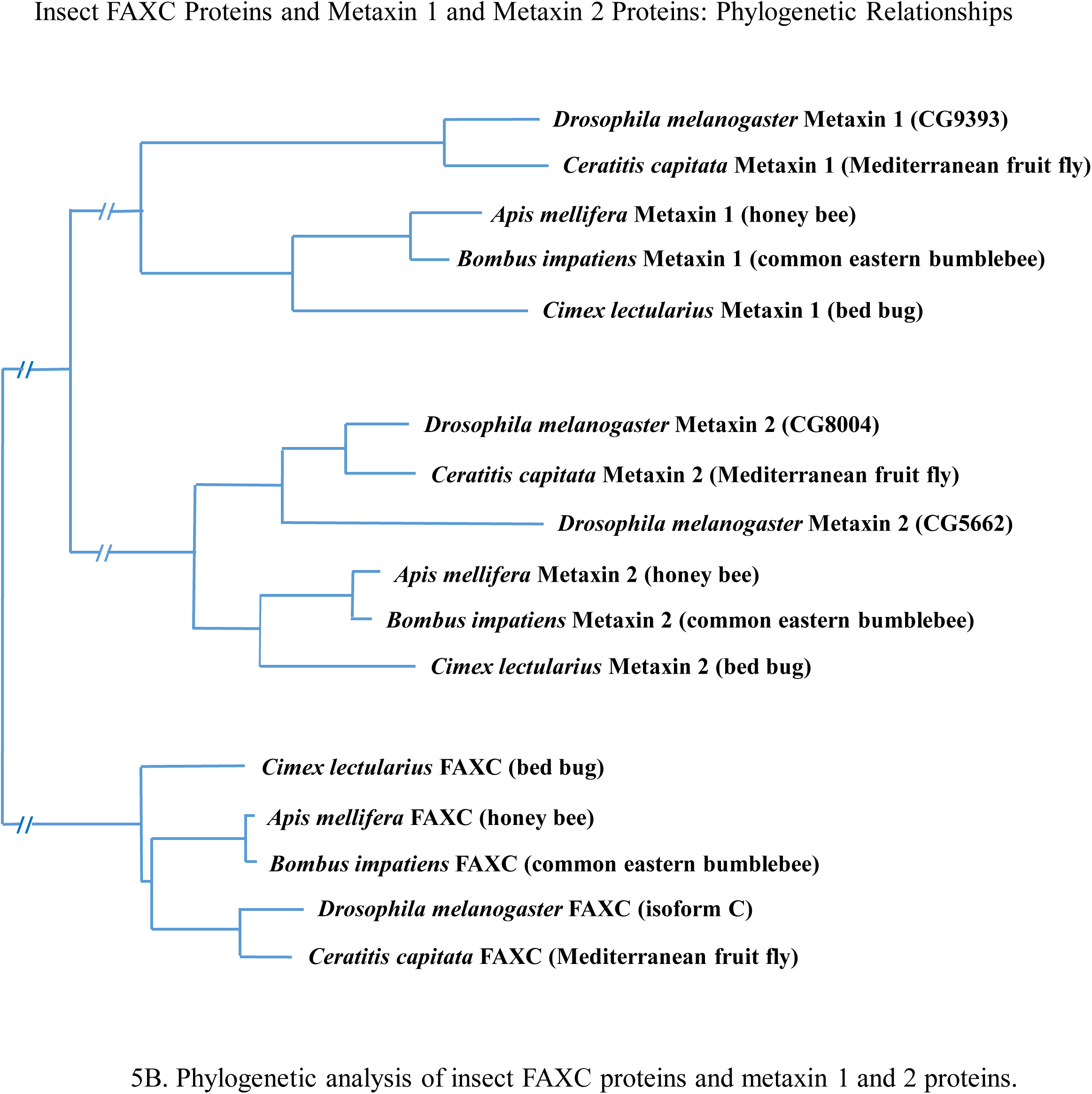
Relationships of invertebrate and vertebrate FAXC and metaxin proteins: phylogenetic analysis. Figure 5A shows that the FAXC proteins of two representative invertebrates are phylogenetically related. FAXC proteins #1 - #4 of both *Aplysia* and *Exaiptasia* form two separate but closely connected groups. These groups are also related to a group of vertebrate FAXC proteins. Similarly, vertebrate metaxin 1 and metaxin 2 proteins form two different but connected groups. Importantly, the FAXC and metaxin proteins seen in Figure 5A are also phylogenetically related to each other. Insect FAXC and metaxin proteins are in Figure 5B. The five different insects represent diverse taxonomic orders – Diptera (*Drosophila*, *Ceratitis*), Hymenoptera (*Apis*, *Bombus*), Hemiptera (*Cimex*) – and form distinct clusters of FAXC, metaxin 1, and metaxin 2 proteins. Within the clusters, similar insects are most closely related, for example *Drosophila melanogaster* (common fruit fly) and *Ceratitis capitata* (Mediterranean fruit fly). The three protein clusters in Figure 5B are separate but related, an important characterstic of FAXC and metaxin proteins.

**Figure 6.**
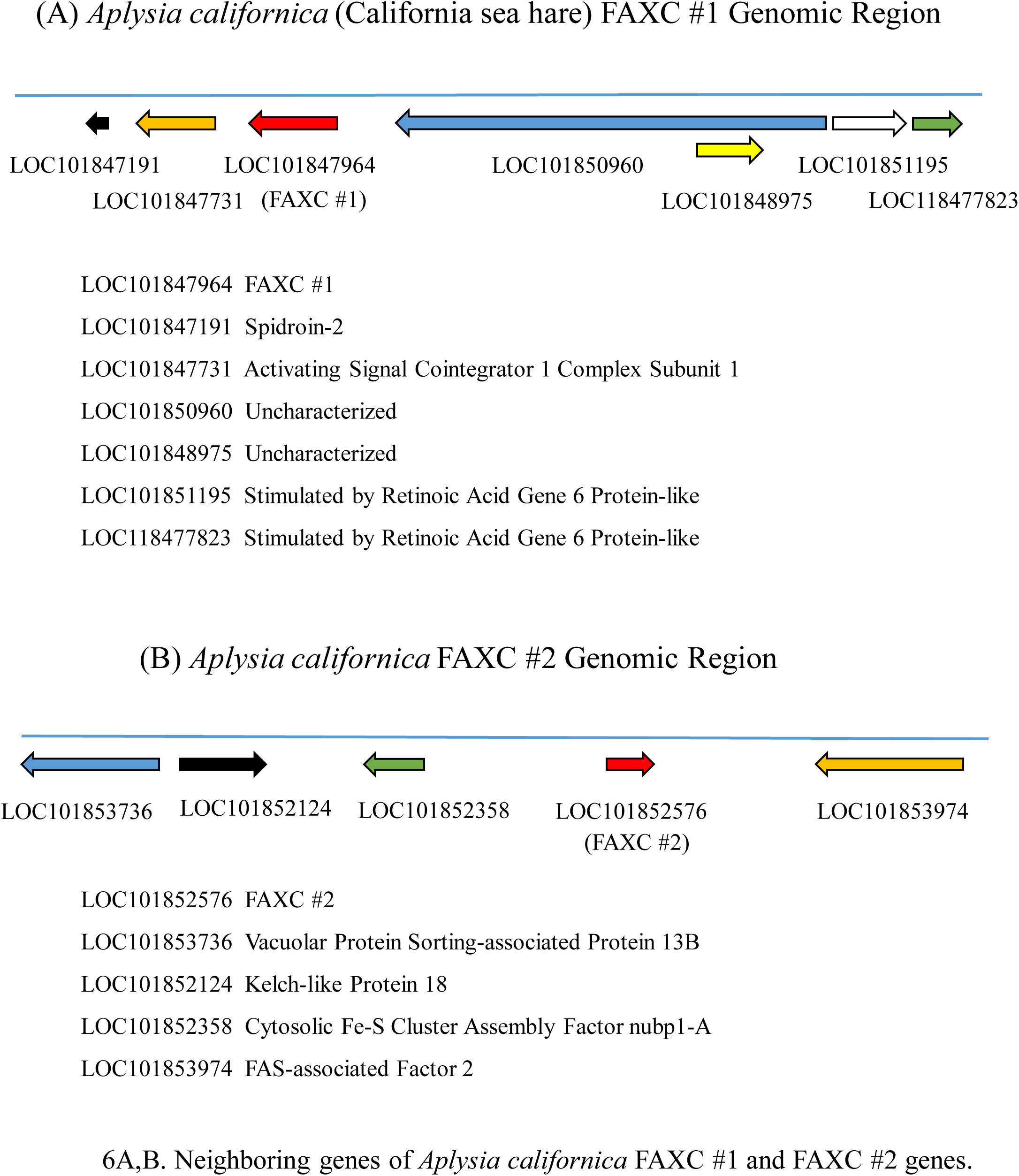
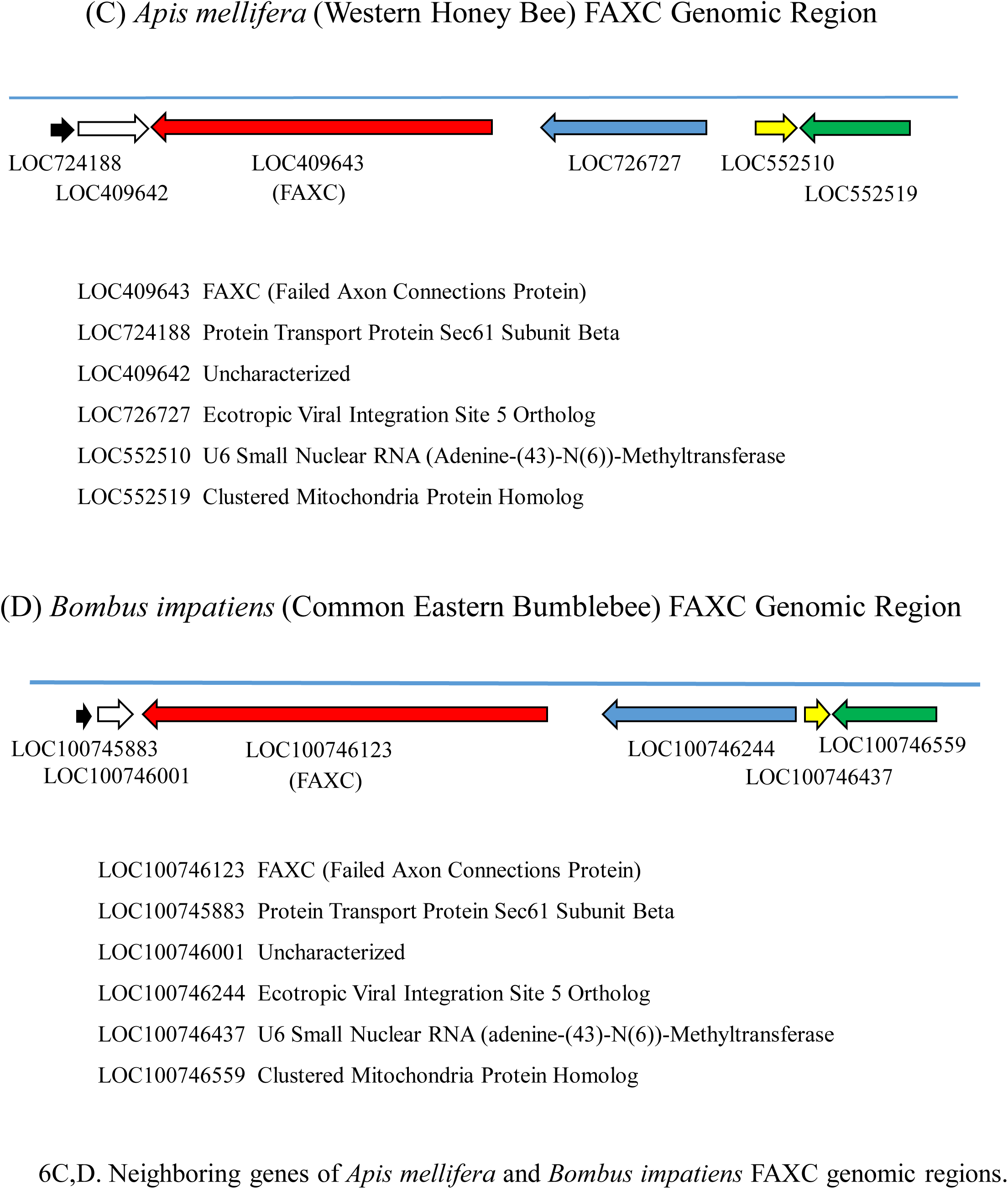
Genomic regions of FAXC genes of invertebrates: neighboring genes. Figure 6A and 6B reveal that the multiple FAXC genes of invertebrates can have different adjacent genes. The examples in 6A and 6B are the FAXC #1 and #2 genes of *Aplysia californica*, an important model invertebrate in neurobiology research. The genes directly adjacent to the two FAXC genes are different, as are the genes that are further from the FAXC genes. In contrast, Figure 6C and 6D show that the neighboring genes can be identical for the FAXC genes of invertebrates that are taxonomically similar. The examples in the figure are the FAXC genes of the honey bee *Apis mellifera* and the bumblebee *Bombus impatiens*. Both are in the Hymenoptera order of Insecta, a large order consisting of bees, wasps, ants, and other species. The neighboring genes of the FAXC genes of the two insects in 6C and 6D are the same, and the genes are in the same order.

**Figure 7.**
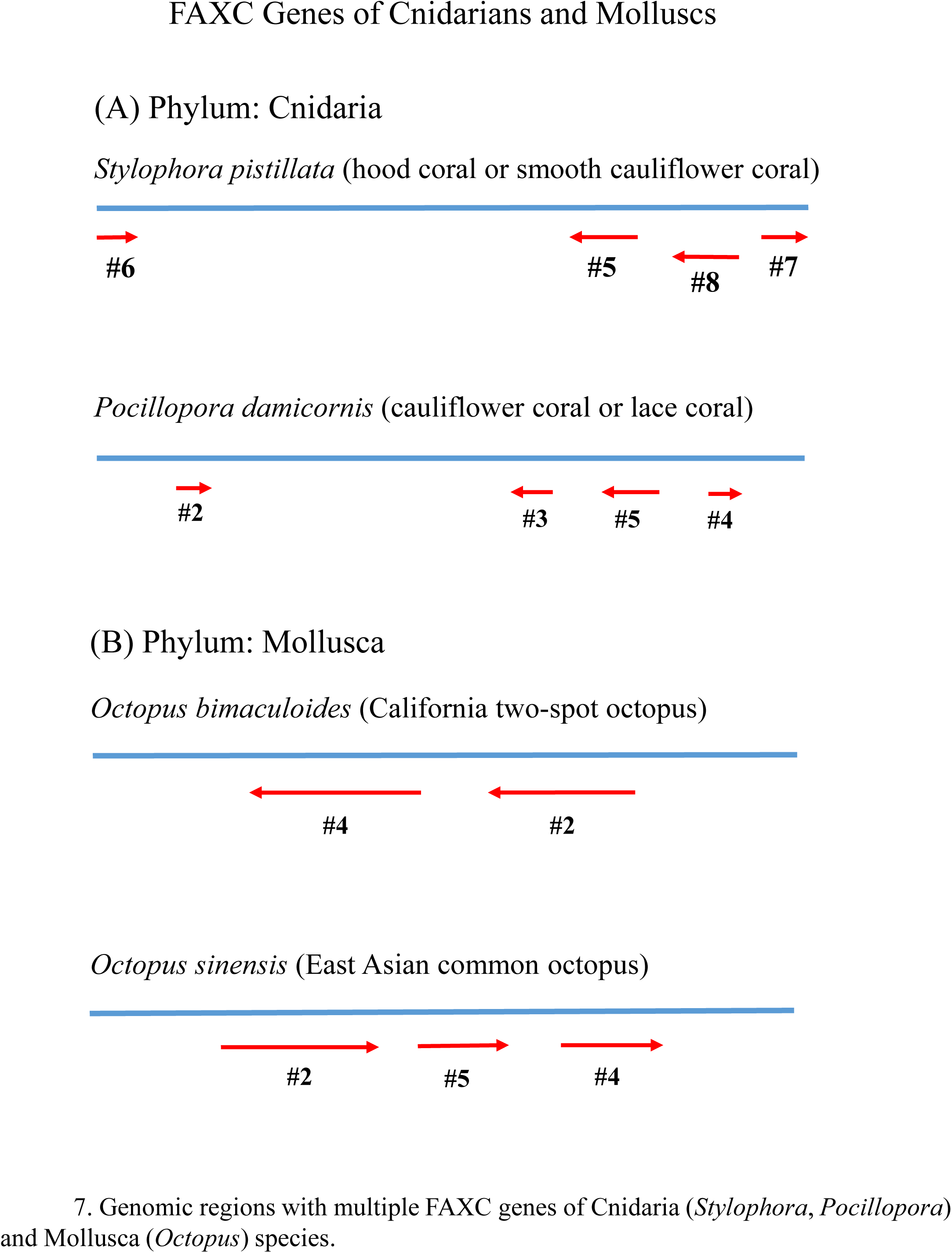
Organization of FAXC genes in genomes of invertebrates with multiple FAXC genes. The figure shows four examples of genomic regions with clusters of FAXC genes for invertebrates that have more than one FAXC gene. The top of the figure includes the genomic region of the coral *Stylophora pistillata* that has four FAXC genes in close proximity. Another coral, *Pocillopora damicornis* (second from top) also has four FAXC genes within the genomic region shown. In addition, the figure includes the FAXC genomic regions of two different octopus species that have multiple FAXC genes. *Octopus bimaculoides* has two FAXC genes in the genomic region in the figure, and *Octopus sinensis* has three. *Stylophora pistillata* has at least nine FAXC genes in total, *Pocillopora damicornis* at least seven, and *O. bimaculoides* and *O. sinensis* each have five or more.

## 3. RESULTS AND DISCUSSION

### 3.1. FAXC and Metaxin Proteins of Invertebrates: Amino Acid Sequence Homology

Pairwise amino acid sequence alignments demonstrate that FAXC and metaxin proteins of invertebrates represent distinct types of proteins. Figure 1A and 1B include alignments of *Branchiostoma floridae* FAXC #1 with *B*. *floridae* metaxin 1 (Figure 1A) and metaxin 2 (Figure 1B). *Branchiostoma floridae*, the Florida lancelet or amphioxus, is an invertebrate of the Chordata phylum. Low percentages of identical amino acids and positives are observed. FAXC #1 and metaxin 1 share only 17% identical amino acids, while FAXC #1 and metaxin 2 have only 19%. Comparable results were found with the FAXC and metaxin proteins of representative invertebrates of other phyla. For example, the mollusc *Octopus bimaculoides*, the California two-spot octopus, showed 17% identical amino acids in aligning FAXC #1 and metaxin 1. For a representative of the Cnidaria phylum, the starlet sea anemone *Nematostella vectensis*, 21% amino acid identities were found in aligning FAXC #3 and metaxin 2. These and other alignments show that FAXC and metaxin proteins are separate categories of proteins, even though they have structural features in common, as discussed in this report.

Invertebrates of a variety of phyla have been found to have genomes that encode multiple FAXC genes. Phyla with species that have more than a single FAXC gene include Cnidaria, Mollusca, Arthropoda, Chordata, Nematoda, and Placozoa. Examples of invertebrates with 10 or more FAXC genes are *Branchiostoma floridae* (Chordata), *Trichoplax adhaerens* (Placozoa), and *Exaiptasia diaphana* (Cnidaria). For some invertebrates, the multiple genes exist as clusters of genes in close proximity.

### 3.2. Conserved GST_N_Metaxin and GST_C_Metaxin Protein Domains of Invertebrate FAXC Proteins

Metaxin proteins are characterized by the presence of GST_N_Metaxin and GST_C_Metaxin protein domains. These domains were first identified as the defining domains of the metaxin proteins of vertebrates, including human, mouse, zebrafish, and *Xenopus*. The domains are major structural features of the three metaxins that have been recognized: metaxins 1, 2, and 3. Two metaxins (metaxins 1 and 2) have been described in invertebrates, and have the same domains as vertebrate metaxins (Adolph, 2020). In addition, metaxin-like proteins of organisms ranging from plants to bacteria possess GST_N_Metaxin and GST_C_Metaxin domains.

FAXC proteins have also been found to contain GST_N_Metaxin and GST_C_Metaxin domains. The proteins were first reported in the fruit fly *Drosophila melanogaster*, and are present in other invertebrates and also in vertebrates. An initial description of structural features of the FAXC proteins of vertebrates and invertebrates has been published (Adolph, 2023). The conserved structural features among FAXC and metaxin proteins include the GST_N_ Metaxin and GST_C_Metaxin protein domains and similar α-helical and β-strand secondary structures. The properties of invertebrate FAXC proteins are further examined in this report. The emphasis is primarily upon the FAXC proteins of three invertebrate phyla: Mollusca, Cnidaria, and Arthropoda.

Figure 2A includes the major protein domains of the FAXC proteins of four representatives of the Mollusca phylum. These include two FAXCs of invertebrates of the Gastropoda class (snails) and two of the Bivalvia class (mussels, oysters). In each case, the GST_N_Metaxin and GST_C_Metaxin domains are the major domains. Also, the Tom37 domain is present in three of the FAXCs. In Figure 2B, three examples of FAXC proteins of invertebrates of the Cnidaria phylum are shown, with two FAXCs of invertebrates in the Hexacorallia class (sea anemones, corals) and one in the Hydrozoa class (hydra). Each is characterized by GST_N_Metaxin and GST_C_Metaxin domains and the Tom37 domain.

The major protein domains of representative arthropods are shown in Figure 2C and 2D. The examples in 2C include two crustaceans, a collembolan arthropod (springtail), and an arachnid. Three insects that are members of three different orders of Insecta are in Figure 2D. As with the molluscan and cnidarian examples in Figure 2A and 2B, GST_N_Metaxin and GST_C_Metaxin domains are the protein domains that dominate the FAXC protein structures.

Tom37 is another protein domain that is a major domain of invertebrate FAXC proteins. The domain is found in some invertebrate FAXCs, but not in all, as Figure 2 demonstrates. Tom37 was first detected in the yeast *Saccharomyces cerevisiae* as part of the SAM protein complex. The complex, in the outer mitochondrial membrane, has a role in the uptake of nascent proteins into mitochondria. The Tom37 domain is widely distributed among FAXC proteins, not only in invertebrates but also in vertebrates including humans.

The protein domain structures of invertebrate FAXCs as seen in Figure 2 are similar to the domain structures of the metaxin proteins of humans, other vertebrates, and invertebrates (Adolph, 2023). This includes all three metaxin proteins that have been identified for vertebrates: metaxin 1, metaxin 2, and metaxin 3. The same domain structure is also found for the human FAXC protein and other vertebrate FAXCs. The presence of GST_N_Metaxin and GST_C_Metaxin domains therefore points out that invertebrate FAXCs are metaxin-like proteins. This is further strengthened by the α-helical and β-strand secondary structures of invertebrate FAXC proteins (Figure 3, discussed below), which show a pattern similar to that of the vertebrate and invertebrate metaxin proteins.

### 3.3. Secondary Structure of Invertebrate FAXC Proteins: Conserved α-Helices

In addition to GST_N_Metaxin and GST_C_Metaxin protein domains, a conserved pattern of α-helical segments is also a defining feature of invertebrate FAXCs, as well as metaxin proteins of invertebrates and vertebrates. This is evident in the examples included in Figure 3. In particular, Figure 3A shows the predicted α-helical segments of the FAXC proteins of representative invertebrates of the Mollusca phylum (first four FAXC sequences) and the Cnidaria phylum (bottom three sequences). The examples in Figure 3A are the same as in Figure 2A,B, and represent a variety of taxonomic classes: for molluscs, the gastropods and bivalves; and for cnidarians, the hexacorallians and hydrozoans.

The α-helix sequences, underlined and colored red, are labeled H1 through H8. The labeling is based on the numbering used for the metaxins, which have a similar pattern of α-helices. The helices in Figure 3A are seen to be highly conserved among all of the FAXC protein chains. The numbers of amino acids between H1 and H2, and between other pairs of α-helices, are similar for the different invertebrates, producing the striking conserved patterns of helices. Also in Figure 3A, S1 through S4 designate the four β-strand segments of the FAXC proteins. The β-strands are near the N-terminal ends and form a planar β-sheet motif characteristic of the FAXCs and metaxins (see section 3.4).

FAXC proteins of a selection of arthropods, along with human metaxin 1, are shown in Figure 3B. The species of arthropods in the figure are the same as in Figure 2C, except for *Homarus americanus*, the American lobster. The arthropods include crustaceans (first three), a *Folsomia* (springtail) species, and an arachnid (*Centruroides*). The same conserved pattern of α-helical segments H1 through H8 is present as in Figure 3A for molluscs and cnidarians. Especially notable is the similarity of the human metaxin 1 pattern of helices compared to the arthropods. β-strand segments S1 – S4 are also included in Figure 3B.

Representatives of the Insecta class of Arthropoda are in Figure 3C. The insects belong to different taxonomic orders: Diptera (*Drosophila melanogaster*), Hymenoptera (*Apis mellifera*), Lepidoptera (*Bombyx mori*), Hemiptera (*Cimex lectularius*), and Coleoptera (*Dendroctonus ponderosae*). Two of the insects (*Bombyx*, *Cimex*) are also in Figure 2D, which shows their conserved protein domains. The FAXC proteins of the insects in Figure 3C display the prominent and distinctive pattern of α-helical segments seen for other invertebrates in Figure 3A and 3B. Helices H1 – H8 in Figure 3C are in almost exact alignment. For clarity, the figure includes only H1 – H8 and not extra helices that are present near the two ends of the protein chains. Figure 3C does include the four conserved β-strand segments S1 – S4 that are near the N-terminus.

### 3.4. Secondary Structure of Invertebrate FAXC Proteins: β-Sheet Motif

As shown in Figure 3, invertebrate FAXC proteins have conserved β-strand segments in addition to α-helices. The β-strand segments are labeled S1 through S4, underlined, and shown with a blue color. The segments are short compared to the α-helical segments, and are typically 4 or 5 amino acids in length. The pattern of β-strands is highly conserved among FAXC proteins of different invertebrate phyla, including Mollusca, Cnidaria, and Arthropoda. As can be seen in Figure 3, β-strand S1 precedes α-helix H1, while S2, S3, and S4 are between H1 and H2. The order is therefore S1 – H1 – S2 – S3 – S4 – H2.

Besides being highly conserved among invertebrate FAXCs, the four β-strands form a three-dimensional structure that is a characteristic feature of invertebrate FAXCs. This 3D β-sheet motif is also found for metaxin proteins of vertebrates and invertebrates, FAXC proteins of vertebrates, and FAXC-like proteins of fungi and bacteria (Adolph, unpublished). Figure 4A,B includes the conserved β-sheet motif found in the FAXC proteins of representative invertebrates of the Mollusca and Cnidaria phyla. The order of the β-strands in three dimensions is S2 – S1 – S3 – S4 in each case. The directions of the segments are ↑ ↑ ↓ ↑. For *Mytilus galloprovincialis* FAXC (Figure 4A), segments S2 and S1 have five amino acids, while S3 and S4 have four. For *Nematostella vectensis* (Figure 4B), all segments have four amino acids.

In three dimensions, the four segments are in close proximity and are present in a nearly planar arrangement, as indicated in the figure. This was observed using the AlphaFold protein structure database (Jumper et al., 2021). The β-sheet motif is also a feature of metaxin proteins, as presented in Figure 4C,D with *Ceratitis capitata* (Mediterranean fruit fly) FAXC in (C) compared to *Ceratitis capitata* metaxin 1 in (D).

### 3.5. FAXC Proteins and Metaxin Proteins of Invertebrates and Vertebrates: Phylogenetic Relationships

The previous sections of this report have demonstrated that FAXC and metaxin proteins of invertebrates and vertebrates share similar protein domain structures and patterns of α-helical and β-strand segments. These similarities are in keeping with the analysis of the phylogenetic relationships between the proteins, as shown in Figure 5. Specifically, Figure 5A includes four FAXC proteins for each of two invertebrates with multiple FAXCs, *Aplysia californica* (Mollusca) and *Exaiptasia diaphana* (Cnidaria). Vertebrate FAXC proteins and metaxin 1 and metaxin 2 proteins (for human, mouse, *Xenopus*, and zebrafish) are also part of the figure. The four *Aplysia* FAXCs and four *Exaiptasia* FAXCs form two separate clusters. The vertebrate FAXCs also form a separate cluster, but a cluster that is phylogenetically related to the invertebrate FAXCs. Metaxin 1 and metaxin 2 proteins of the same vertebrates form two separate clusters. The vertebrate metaxin 1 and 2 proteins are seen to be less related to the vertebrate and invertebrate FAXCs than to each other. But all of the proteins, FAXCs and metaxins, are shown to be phylogenetically related.

Figure 5B examines the phylogenetic relationships between FAXC proteins and metaxin proteins of insects (Phylum: Arthropoda; Class: Insecta). The figure shows three groups of proteins: metaxin 1, metaxin 2, and FAXC proteins. Invertebrates including insects have metaxin 2 proteins in addition to metaxin 1 proteins, although metaxin 3 proteins have not been detected in invertebrates. Figure 5B demonstrates that insect FAXC and metaxin 1 and 2 proteins are all phylogenetically related. As with the invertebrates of the Mollusca and Cnidaria phyla in Figure 5A, the metaxin 1 and 2 groups are more closely related to each other than to the FAXCs. Within the groups, the closest relationships are between the FAXCs or metaxins of the same taxonomic order. In particular, *Drosophila melanogaster* (common fruit fly) and *Ceratitis capitata* (Mediterranean fruit fly) are both Diptera and are closely related, as are *Apis mellifera* (western honey bee) and *Bombus impatiens* (common eastern bumblebee), which are both Hymenoptera.

### 3.6. Gene Neighbors of Invertebrate FAXC Genes

Invertebrates can have multiple FAXC genes, and the question arises whether the genomic regions of the different FAXC genes are similar, that is whether the adjacent genes are the same or almost the same. The answer to this question can be observed in the examples in Figure 6A,B, which compares the genomic regions of *Aplysia californica* FAXC #1 and FAXC #2. *Aplysia californica*, the California sea hare or sea slug, is an important laboratory animal used in neurobiology studies. In Figure 6A and 6B, the neighboring genes of FAXC #1 and FAXC #2 are seen to be different. The FAXC #1 gene is between the gene for activating signal cointegrator 1 complex subunit 1 and an uncharacterized gene. In contrast, the FAXC #2 gene is between a gene for FAS-associated factor 2 and a gene for cytosolic Fe-S cluster assembly factor nubp1-A. Other neighboring genes are also different.

However, invertebrates that are taxonomically similar can have similar genomic regions. This is demonstrated in Figure 6C,D, which compares the adjacent genes for the western honey bee *Apis mellifera* and the common eastern bumblebee *Bombus impatiens*. The genes are seen to be identical and in the same order. For the honey bee (Figure 6C), the FAXC gene is flanked on one side by the gene for protein transport protein Sec61 subunit beta and an uncharacterized gene, and on the other side by the gene for ecotropic viral integration site 5 ortholog. For the bumblebee (Figure 6D), the genes in the genomic region shown are identical to the honey bee genes.

### 3.7. Genomic Regions of Invertebrates with Multiple FAXC Genes

As the previous sections of this report have indicated, invertebrates can have more than one FAXC gene, and in some cases many more. Species with multiple FAXC genes can be found in phyla that include Cnidaria, Mollusca, Arthropoda, Chordata, and Placozoa. Figure 7 shows genomic regions with multiple FAXC genes for invertebrates from two phyla: Cnidaria (*Stylophora* and *Pocillopora*) and Mollusca (two Octopus species). *Stylophora* has at least 9 FAXC genes, *Pocillopora* at least 7, and the two *Octopus* species at least 5 each.

The multiple FAXC genes are frequently found as groups of genes in close proximity in a genomic region, as Figure 7 reveals. For example, four of the FAXC genes of the coral *Stylophora pistillata* (#6, #5, #8, #7) are in close proximity in the genomic region included in Figure 7 (top). Another coral of the Cnidaria phylum, *Pocillopora damicornis*, also has four FAXC genes that form a cluster of adjacent genes. And the two *Octopus* species of the Mollusca phylum each have two or three FAXC genes close together.

In contrast to the situation with invertebrates, a single FAXC gene has generally been detected in vertebrates. With the vertebrates included in this report (human, mouse, *Xenopus*, zebrafish), human and mouse each have one FAXC gene, although with isoforms. But, due to genome duplication, both *Xenopus* and zebrafish have two FAXC genes. In regard to metaxin genes, vertebrates possess genes for three metaxin proteins: metaxin 1, metaxin 2, and metaxin 3. For invertebrates, two metaxin genes, homologous to vertebrate metaxins 1 and 2, have been identified. But the situation can be more complicated. *Drosophila melanogaster*, for example, has one gene for metaxin 1, but two genes for metaxin 2.

